# Scp160 deletion suppresses TDP-43 aggregation and toxicity in *Saccharomyces cerevisiae*

**DOI:** 10.64898/2026.04.28.721091

**Authors:** Joan Castells-Ballester, Natalia Shcherbik

## Abstract

Protein misfolding and aggregation are central features of neurodegenerative disease, yet the cellular factors that determine whether aggregation-prone proteins become toxic remain incompletely understood. The RNA-binding protein TDP-43 forms cytoplasmic aggregates in amyotrophic lateral sclerosis (ALS) and is toxic when expressed in yeast, providing a tractable model to identify modifiers of TDP-43 proteotoxicity. Scp160 and Bfr1 are yeast RNA-binding/translation-associated proteins linked to polysomes, ER-localized transcripts, and mRNP organization, but their contribution to TDP-43 aggregation has not been tested. Here, we expressed human TDP43-GFP in *Saccharomyces cerevisiae* strains lacking *SCP160* or *BFR1* and examined TDP43-GFP abundance, toxicity, and aggregate formation. Although deletion of *SCP160* or *BFR1* did not cause major changes in bulk translation, *scp160Δ* cells accumulated higher levels of TDP43-GFP protein, yet showed improved growth during prolonged induction and a marked reduction in severe cytoplasmic aggregates. Loss of *BFR1* produced a more intermediate phenotype, reducing total aggregate-positive cells but showing a weaker effect on severe aggregates and toxicity. Thus, deletion of *SCP160* uncouples TDP43-GFP abundance from toxicity and severe aggregate formation. These findings identify Scp160 and, to a lesser extent, Bfr1 as yeast factors that promote visible TDP43-GFP aggregate formation and toxicity independently of reduced protein abundance.

## Introduction

Protein misfolding and aggregation are central pathogenic features of multiple neurodegenerative disorders, including Alzheimer’s disease, Huntington’s disease, Parkinson’s disease, and amyotrophic lateral sclerosis (ALS). In these proteinopathies, misfolded proteins accumulate as soluble oligomers, insoluble aggregates, or other aberrant species that disrupt cellular homeostasis, engage in aberrant interactions, and overload the proteostasis machinery, ultimately contributing to cellular stress, dysfunction, and death (reviewed in [1, 2]). In this context, the unicellular eukaryote *Saccharomyces cerevisiae* has proven to be a powerful model for studying protein-misfolding disorders because of its conserved protein quality control pathways and its tractability for genetic analysis. Yeast models have successfully recapitulated key aspects of aggregation and toxicity for several disease-associated proteins, including α-synuclein, polyalanine-expanded proteins, TDP-43, and FUS [3–10].

A pathological hallmark of ALS is the abnormal accumulation and aggregation of the RNA-binding protein TDP-43. TDP-43 is a predominantly nuclear RNA-binding protein that regulates multiple aspects of RNA metabolism [11, 12]. Under pathological conditions, however, TDP-43 can mislocalize to the cytoplasm, where it becomes prone to aggregation and contributes to cellular toxicity and neurodegeneration [11, 13] [14]. In addition to protein aggregation itself, TDP-43 proteinopathy is closely linked to disturbed RNA metabolism, translational control, and cytoplasmic ribonucleoprotein (RNP) granule dynamics (reviewed in [14, 15]). Yeast TDP-43 models reproduce key features of this pathology, including cytoplasmic accumulation and toxicity, and have proven useful for identifying genetic modifiers of TDP-43-driven proteotoxicity [16–20]. Unlike polyglutamine proteins, whose aggregation is largely driven by expansion of a defined glutamine-rich tract, TDP-43 aggregation reflects a more complex pathogenic process involving cytoplasmic mislocalization, a prion-like low-complexity C-terminal domain, and disturbed RNA-dependent RNP dynamics.

Among yeast RNA-binding proteins, Scp160 emerged as a particularly plausible candidate modifier of TDP-43 proteotoxicity. Previous work implicated Scp160 in regulating aggregation-prone proteins in yeast, showing that both the aggregation and toxicity of exogenous polyQ reporters are influenced by Scp160, and that Scp160 also affects the aggregation of endogenous Q/N-rich proteins. In addition, Scp160 is a multi-KH-domain RNA-binding protein associated with polysomes and ER-localized transcripts, where it contributes to mRNA selection and translational efficiency [21–23]. Bfr1, a functionally linked partner of Scp160, is likewise a polysome-associated protein implicated in the translational regulation of selected transcripts, including secretory-pathway proteins [24–26]. In addition, both Scp160 and Bfr1 have been connected to the regulation of P-body dynamics under normal growth conditions [27–29]. Together, these properties place Scp160 and Bfr1 at the intersection of RNA handling, selective translation, and cytoplasmic mRNP organization, raising the possibility that they may influence TDP-43 proteotoxicity.

Yeast TDP-43 models have identified multiple modifiers of aggregation and toxicity, but the contribution of ER-associated RNA-binding and translation regulators such as Scp160 has not been tested. Here, we used a *S. cerevisiae* TDP-43 model to test whether loss of Scp160 or Bfr1 alters TDP-43 abundance, aggregation, and toxicity. By defining these phenotypes in a genetically tractable yeast system, this study provides an initial framework for evaluating whether ER-associated RNA-binding and translation regulators influence TDP-43 proteotoxicity.

## Results

### Bulk translation is preserved in scp160Δ and bfr1Δ cells

Scp160 has previously been characterized as a polysome-associated RNA-binding protein that regulates translation of selected mRNA subsets without causing overt changes in bulk translation upon acute depletion [22]. As Hirschmann et al.’s study used a W303 background, wherein SCP160 was under the control of the Dox-repressible promoter, resulting in depletion of Scp160 over time [22], while our experimental system employed a viable *scp160Δ* strain on BY4742 background, we first examined polysome profiles to determine whether the complete loss of Scp160 would affect global translation. Because Bfr1 is a functionally linked partner of Scp160, and *BFR1* deletion has been reported to alter Scp160 distribution across sucrose-gradient fractions [24], *bfr1Δ* cells were analyzed in parallel. Cell lysates were prepared from mid-log cultures of *WT, scp160Δ*, and *bfr1Δ* cells grown in YPDA medium and analyzed by sucrose density centrifugation. Consistent with published data [22, 24, 26, 29], ribosomal species sedimentation profiles of *scp160Δ*- and *bfr1Δ*-derived lysates were similar to those of *WT*, with no striking changes in the relative abundance of free subunits, 80S monosomes, or polysomes (Fig. 1A). Thus, loss of *SCP160* or *BFR1* did not reveal an obvious defect in bulk translation in the BY4742 background, consistent with prior studies and supporting the suitability of these strains for analysis of TDP-43-associated phenotypes.

**Figure 1.**
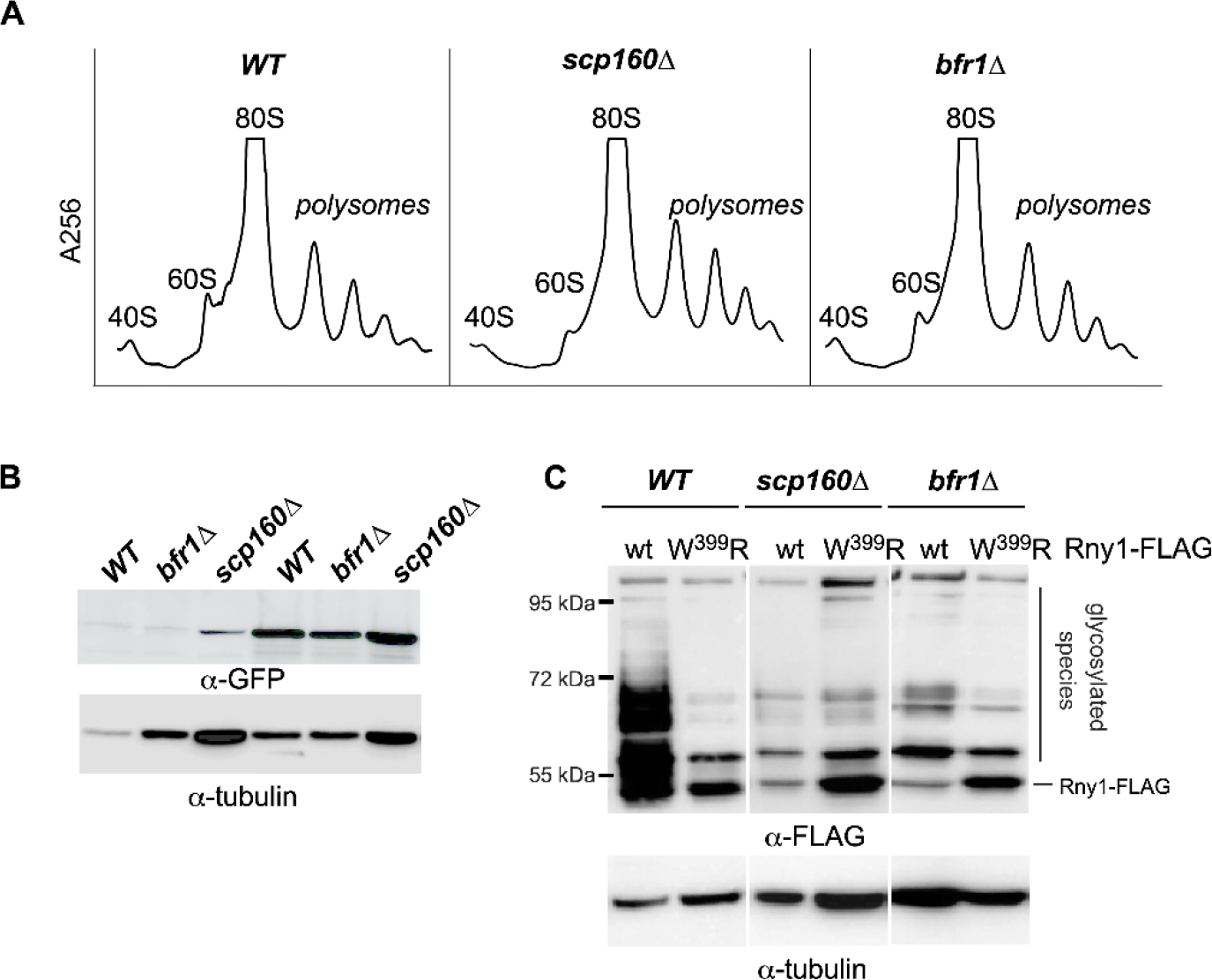
Loss of *SCP160* or *BFR1* does not cause major changes in bulk ribosome sedimentation or TDP43-GFP accumulation. **(A)** Ribosome sedimentation profiles of *WT, scp160Δ*, and *bfr1Δ* cells. Cells were grown in YPDA to mid-log phase and lysed under polysome-preserving conditions. Lysates were centrifuged through 15-45% sucrose gradients and fractionated with the continuous measurement of absorbance at 256 nm to visualize ribosomal species peaks. Positions of 40S and 60S ribosomal subunits, 80S monosomes, and polysomes are indicated. **(B)** Semi-quantitative immunoblot analysis of TDP43-GFP expression in *WT, bfr1Δ*, and *scp160Δ* cells carrying *P*_*GAL1*_-TDP43-GFP. Cells were grown on raffinose and then shifted to either glucose-repressing conditions or to galactose for induction for 4 h. TDP43-GFP was detected with an anti-GFP antibody; tubulin was used as a loading control. **(C)** Immunoblot analysis of secretory-protein reporter Rny1-FLAG in *WT, scp160Δ*, and *bfr1Δ* cells. Rny1-FLAG (wild-type (wt) or the glycosylation-defective Rny1-FLAG W399R mutant) was expressed from the 2 m plasmid from the *ADH* promoter. Rny1-FLAG species were detected with anti-FLAG antibody. Tubulin was used as a loading control.

### TDP43-GFP accumulates to comparable levels in WT, bfr1Δ, and scp160Δ cells

To test whether Scp160 and Bfr1 influence TDP-43 behavior in the BY4742 background of *Saccharomyces cerevisiae*, we expressed human TDP43-GFP from a 2μ plasmid under control of the GAL1 promoter [6] in *WT, scp160Δ*, and *bfr1Δ* cells. This inducible system allowed us to avoid constitutive expression of toxic TDP-43 and to monitor its accumulation after a shift from glucose-to galactose-containing medium.

We first asked whether deletion of *SCP160* or *BFR1* affects steady-state TDP43-GFP protein levels. Cells were grown in raffinose-containing medium, shifted to glucose or galactose, and collected after 4 h of induction. As expected, no TDP43-GFP signal was detected by western blotting using anti-GFP antibodies in glucose-grown cultures, consistent with repression of the GAL1 promoter, whereas a clear signal was observed after galactose induction in all three strains. Anti-tubulin immunoblotting was included as a loading control to verify lysate loading across lanes. Because *SCP160* and *BFR1* have been implicated in selective translational regulation, tubulin was not used here as a formal normalization factor for quantitative comparison among strains. Accordingly, this analysis was interpreted as semi-quantitative and used to assess whether any major genotype-dependent differences in TDP43-GFP accumulation were evident. Under these conditions, no major difference in TDP43-GFP abundance was apparent among *WT, bfr1Δ*, and *scp160Δ* cells 4 h post-galactose induction (Fig. 1B).

In contrast, Rny1-FLAG provided a yeast secretory-protein comparison for assessing Scp160/Bfr1-sensitive expression behavior in the same strain background. Rny1 was selected because *RNY1* is a reported Bfr1-associated transcript whose expression is influenced by Bfr1-dependent post-transcriptional regulation [25, 30]. In previous work from our laboratory, these Rny1-FLAG constructs were used to define the secretory pathway processing of Rny1, including the Golgi glycosylation-defective W399R mutant [31]. Here, we used these previously characterized Rny1-FLAG constructs as a yeast protein comparison for protein-expression changes in *WT, scp160Δ*, and *bfr1Δ* cells. As expected, wild-type Rny1-FLAG migrated as multiple species, consistent with secretory-pathway processing, whereas W399R lacked the heterogeneous high-molecular-weight glycosylated forms characteristic of Golgi-dependent modification (Fig. 1C).

Across the strain comparison, the major Rny1-FLAG species were preserved, indicating that deletion of *SCP160* or *BFR1* did not grossly alter Rny1 secretory-pathway processing. However, the overall Rny1-FLAG signal differed between strains, consistent with altered Rny1 protein expression in the absence of *BFR1* and, similarly, in the absence of its functional partner *SCP160*. This expected Rny1-FLAG response supported the reliability of the strain comparison used to evaluate TDP43-GFP expression. Importantly, TDP43-GFP did not exhibit genotype-dependent changes in abundance, suggesting that the reduced TDP-43 toxicity and aggregation observed in *scp160Δ* and *bfr1Δ* cells are unlikely to result from reduced TDP43-GFP expression.

### Galactose induction leads to elevated TDP43-GFP fluorescence in scp160Δ cells

To corroborate the immunoblot analysis with an independent live-cell approach, we monitored TDP43-GFP fluorescence in cultures following induction from the GAL1 promoter while simultaneously recording OD600 as an estimate of culture density. This assay also avoids potential artifacts associated with cell lysis and partial recovery of insoluble material during immunoblot preparation. *WT, bfr1Δ*, and *scp160Δ* cells carrying *P*_*GAL1*_ TDP43-GFP were grown in raffinose medium to mid-log phase and shifted to glucose- or galactose-containing medium. As expected, glucose cultures displayed only background fluorescence, whereas galactose induced progressive accumulation of GFP signal in all three strains (Fig. 2A). The increase was most pronounced in *scp160Δ* cells.

**Figure 2.**
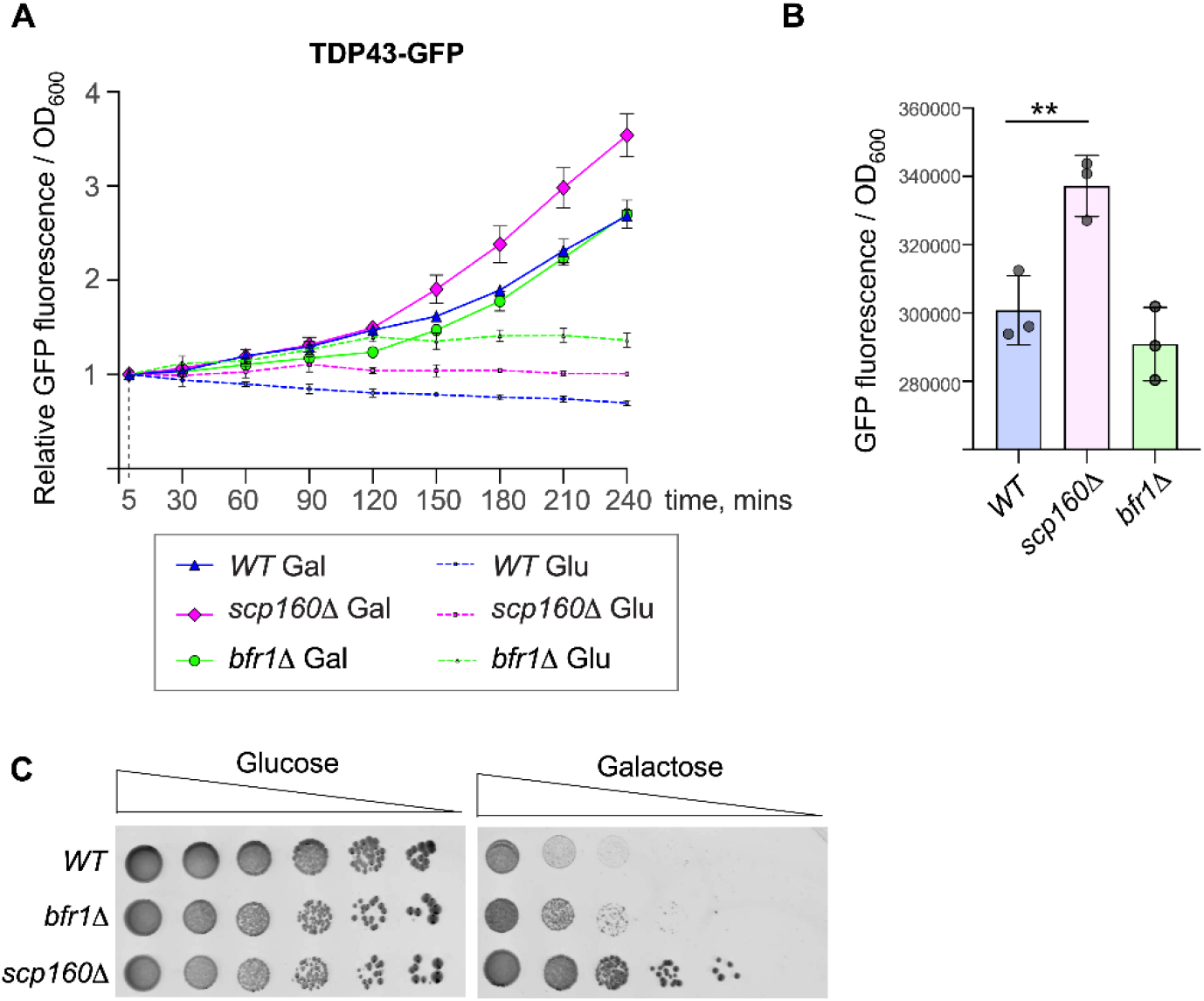
Loss of *SCP160* reduces TDP43-GFP toxicity despite increased normalized fluorescence signal. **(A)** Time-course analysis of TDP43-GFP fluorescence in live cultures of *WT, bfr1Δ*, and *scp160Δ* cells carrying *PGAL1*-TDP43-GFP. Cells were grown overnight in synthetic raffinose medium lacking uracil, diluted into fresh medium, and cultured to mid-log phase (OD600 ≈ 0.5). Cultures were then shifted to synthetic medium containing either glucose (repressing) or galactose (inducing), and GFP fluorescence and OD at l=600 nm were measured at the time points indicated in the Figure using a BioTek Synergy plate reader. Fluorescence values were normalized to culture density (OD_600_) and expressed relative to the first time point. Data represent mean ± SD. **(B)** Quantification of normalized TDP43-GFP fluorescence after 4 h of galactose induction. GFP fluorescence from the same cultures shown in (A) was divided by OD_600_. Bars represent mean values; individual biological replicates are shown as symbols (*n* = 3). Statistical analysis was performed using ordinary one-way ANOVA followed by Dunnett’s multiple-comparisons test using *WT* as the control group. Overall ANOVA: F(2,6) = 17.93, *P* = 0.0029. Dunnett’s test: *WT* vs. *scp160Δ*, adjusted *P* = 0.0076 (**); *WT* vs. *bfr1Δ*, adjusted *P* = 0.4230 (ns). **(C)** Serial dilution growth assay demonstrating the effect of prolonged TDP43-GFP expression on cell fitness. *WT, bfr1Δ*, and *scp160Δ* strains carrying *P*_*GAL1*_-TDP43-GFP were grown overnight at 30 °C in synthetic raffinose medium lacking uracil, diluted to OD600 ≈ 0.2, and cultured for an additional 4 h. Cells were washed, adjusted to 2 × 10^6^ cells/ml, and subjected to six 1:5 serial dilutions. Dilutions were spotted onto synthetic agar containing glucose (repressing) or galactose (inducing) as the carbon source. Plates were incubated at 30°C for 7 days before imaging.

Normalization of endpoint fluorescence values to OD_600_ after 4 h of induction revealed significantly higher TDP43-GFP fluorescence in *scp160Δ* cultures relative to *WT* cells, whereas *bfr1Δ* cells were not significantly different from *WT* (Fig. 2B). These data are consistent with the immunoblot trend (Fig. 1B) and indicate that loss of *SCP160* does not reduce but rather increase the TDP43-GFP abundance. Instead, *scp160Δ* cells accumulate equal or greater levels of the protein despite exhibiting reduced toxicity (see Fig. 2C), suggesting that suppression is mediated by altered handling of TDP-43 rather than decreased expression.

### Deletion of SCP160 suppresses TDP43-GFP toxicity during prolonged induction

Since the fluorescence and immunoblot experiments were performed after a relatively short induction period, we next asked whether sustained TDP43-GFP expression produced different longer-term effects on cell fitness. Equal numbers of *WT, bfr1Δ*, and *scp160Δ* cells carrying *P*_*GAL1*_-TDP43-GFP were spotted onto glucose- or galactose-containing agar plates. All three strains grew similarly on glucose, whereas galactose-induced TDP43-GFP expression strongly impaired the growth of *WT* cells. In contrast, deletion of *SCP160* markedly improved growth under inducing conditions, indicating partial suppression of TDP43-dependent toxicity. Deletion of BFR1 produced a weaker rescue phenotype under inducing conditions (Fig. 2C). Thus, despite accumulating equal or greater levels of TDP43-GFP, *scp160Δ* cells were less sensitive to TDP43 toxicity, suggesting that loss of *SCP160* alters the cellular handling of TDP43 rather than simply reducing its expression.

### Deletion of SCP160 preferentially reduces severe TDP43-GFP aggregation

We next analyzed TDP43-GFP localization by live fluorescence microscopy after 4 h of galactose induction. In *WT* cells, TDP43-GFP was detected predominantly in the cytoplasm and frequently appeared in bright punctate foci, consistent with the aggregation-prone behavior of TDP-43 in yeast models [17]. DAPI staining confirmed that these GFP-positive structures were extranuclear (Fig. 3A).

**Figure 3.**
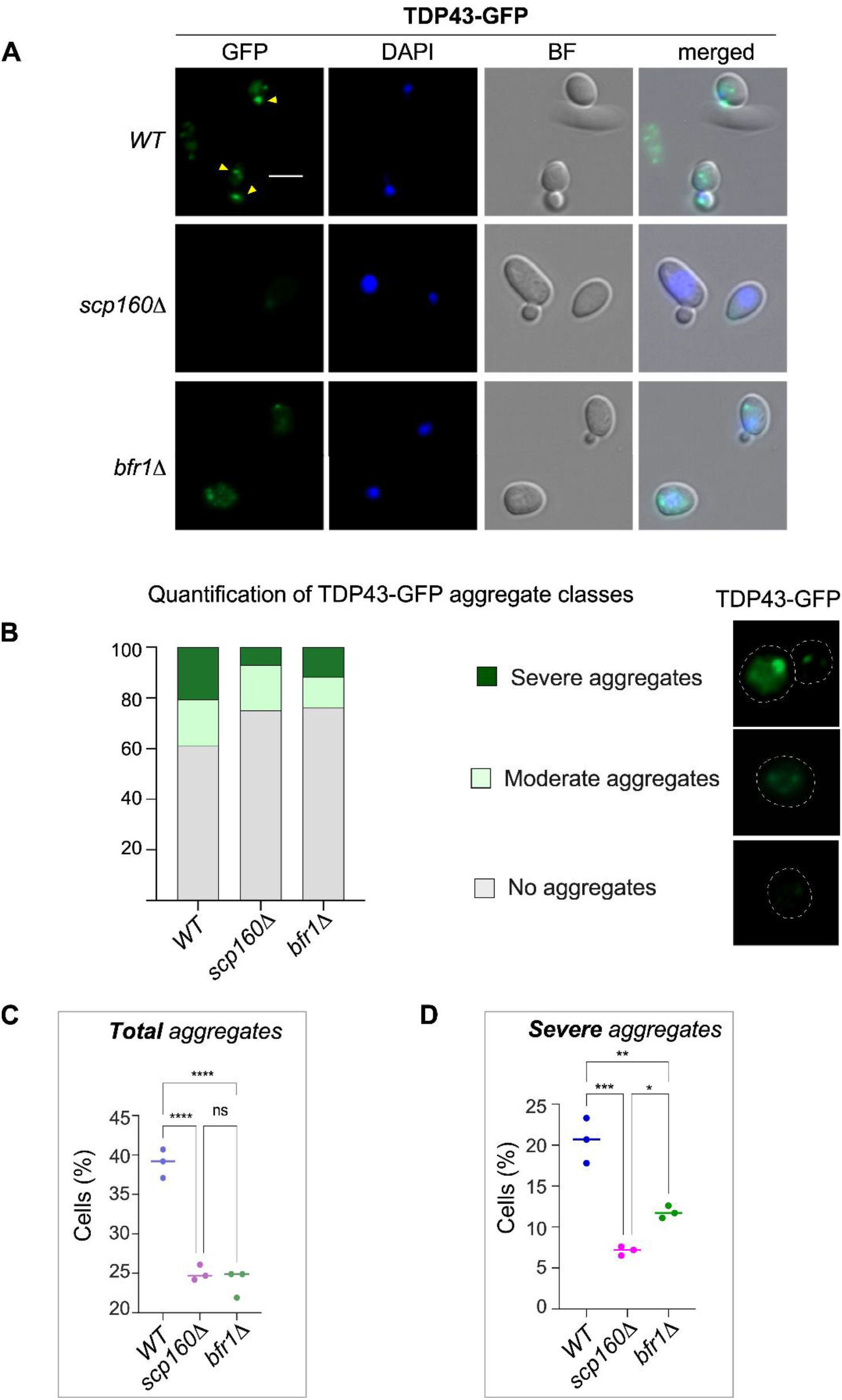
Loss of *SCP160* and *BFR1* reduces TDP43-GFP aggregation, with *SCP160* deletion producing the strongest reduction in severe aggregates. **(A)** Representative fluorescence microscopy images of *WT, scp160Δ*, and *bfr1Δ* cells expressing *P*_*GAL1*_-TDP43-GFP. Cells were grown in synthetic raffinose medium lacking uracil to mid-log phase, shifted to galactose-containing medium for 4 h to induce TDP43-GFP expression, and stained with DAPI to visualize nuclei. GFP, DAPI, bright-field (BF), and merged images are shown. Yellow arrowheads indicate bright cytoplasmic TDP43-GFP foci. Scale bar, 10 µm. **(B)** Quantification of TDP43-GFP aggregate classes in *WT, scp160Δ*, and *bfr1Δ* cells after 4 h of galactose induction. Cells were classified as having no visible aggregates, moderate aggregates, or severe aggregates based on GFP signal pattern. Moderate aggregates were defined as one or more small puncta, whereas severe aggregates were defined as large single foci and/or multiple prominent puncta within the same cell. The experiment was performed in triplicate, with 100-200 cells counted per strain in each replicate. Representative examples of each aggregate class are shown on the right. Dashed white outlines indicate cell boundaries. **(C)** Quantification of total aggregate-positive cells from the data shown in Fig. 3B. Total aggregate-positive cells were defined as cells containing either moderate or severe TDP43-GFP aggregates. Dots represent independent experiments, and horizontal lines indicate the mean. Statistical significance was determined using ordinary one-way ANOVA followed by Tukey’s multiple comparisons test. *WT* vs. *scp160Δ, P* < 0.0001 (****); *WT* vs. *bfr1Δ, P* < 0.0001 (****); *scp160Δ* vs. *bfr1Δ, P* = 0.6786 (non-significant, ns). **(D)** Quantification of cells with severe TDP43-GFP aggregates from the data shown in Fig. 3B. Dots represent independent experiments, and horizontal lines indicate the mean. Statistical significance was determined using ordinary one-way ANOVA followed by Tukey’s multiple comparisons test. *WT* vs. *scp160Δ, P* = 0.0002 (***); *WT* vs. *bfr1Δ, P* = 0.0016 (**); *scp160Δ* vs. *bfr1Δ, P* = 0.0323 (*).

Visual inspection revealed clear strain-dependent differences: *WT* cells commonly contained bright cytoplasmic TDP43-GFP foci, whereas *bfr1Δ* cells still formed visible foci, but these were consistently smaller and less intense. In contrast, *scp160Δ* cells displayed a marked reduction in visible TDP43-GFP foci, with many cells showing diffuse, weak, or no obvious aggregate-like signal (Fig. 3A).

To quantify these differences, cells were classified into three categories: no visible aggregates, moderate aggregates, or severe aggregates (Fig. 3B). In *WT* cultures, approximately 39% of cells contained visible TDP43-GFP aggregates, with moderate and severe aggregates each representing about 20% of the population. Deletion of either *SCP160* or *BFR1* significantly reduced the total fraction of aggregate-positive cells compared to *WT* (Fig. 3C). Notably, total aggregate levels were not significantly different between *scp160Δ* and *bfr1Δ* cells, indicating that both deletions reduced the overall frequency of visible TDP43-GFP aggregation to a similar extent.

However, analysis of aggregate severity revealed a clear distinction between the two mutants. Severe aggregates were strongly reduced in *scp160Δ* cells compared to either *WT* or *bfr1Δ* cells (Fig. 3D). By comparison, *bfr1Δ* cells showed an intermediate phenotype, with fewer severe aggregates than *WT* but more severe aggregates than *scp160Δ*. Thus, although deletion of either *SCP160* or *BFR1* reduced the total aggregate-positive population (Fig. 3C), loss of *SCP160* produced the strongest shift away from the severe aggregate phenotype (Fig. 3D).

These differences in aggregate burden and severity are consistent with the degree of toxicity rescue observed in each mutant strain (Fig. 2C) and with prior yeast studies associating TDP-43 aggregation state with cellular toxicity [16]. Together, these data suggest that Scp160 and, to a lesser extent, Bfr1 promote the formation or maintenance of visible TDP43-GFP aggregates in yeast.

## Discussion

Yeast models have been highly informative for dissecting TDP-43 aggregation and toxicity [16, 17]. Prior work showed that TDP-43 forms cytoplasmic foci in yeast and that aggregation-prone variants are associated with increased toxicity [16]. Genetic screens have also identified RNA-processing and proteostasis pathways as modifiers of TDP-43 toxicity [17, 32]. Our findings extend this framework by revealing an unexpected relationship between TDP43-GFP abundance, aggregation, and toxicity in yeast cells lacking the RNA-binding proteins Scp160 or Bfr1.

The increased TDP43-GFP abundance in *scp160Δ* cells (Fig. 2B) argues against a simple model in which toxicity is driven solely by protein overproduction or insufficient folding capacity. If TDP43-GFP toxicity were caused only by excessive protein accumulation overwhelming the folding machinery, then *scp160Δ* cells would be expected to exhibit equal or greater toxicity. Instead, cells lacking *SCP160*, and to a lesser extent *BFR1*, accumulated at least as much TDP43-GFP protein as *WT* cells (Figs. 1B and 2B), but displayed improved growth and fewer severe aggregates (Figs. 2C and 3). These observations support a model in which Scp160 influences the translation-associated handling of TDP43-GFP, thereby affecting whether newly synthesized protein enters a severe aggregate-prone state.

This model is consistent with prior work showing that Scp160p is not required for global polysome formation but supports efficient translation of selected mRNAs [22]. Hirschmann et al. showed that depletion of Scp160p did not substantially alter global polysome-to-monosome ratios but altered the translational state of a subset of mRNAs, many of which encode cell wall, extracellular, or ER/Golgi-related proteins [22]. Importantly, for the Scp160p target *PRY3*, a shift toward heavier polysome fractions was accompanied by reduced protein output, which the authors interpreted as “slow polysomes” caused by impaired elongation rather than increased translation [22]. They further proposed that Scp160p may promote translational efficiency through mechanisms related to codon order, tRNA reuse, and prevention of tRNA diffusion from translating ribosomes [22].

Although TDP43-GFP is not a native Scp160 target mRNA, it is a heterologous protein expressed from a strong inducible promoter *GAL1* [33], and its folding or partitioning may be highly sensitive to the local translation environment. Loss of *SCP160* could alter elongation dynamics, ribosome-associated factor recruitment, tRNA availability, or the timing of cotranslational folding events [22, 34, 35]. Such changes could redirect TDP43-GFP away from the formation or maintenance of large visible aggregates without reducing total protein accumulation [36].

Asc1/RACK1 provides a possible link between Scp160/Bfr1-associated mRNPs and ribosome-based regulation. Scp160p has been reported to associate with the 40S subunit near the mRNA-binding site, and this ribosome association is partially dependent on Asc1p/RACK1 [22, 27]. Asc1 is a conserved 40S ribosomal scaffold implicated in selective translational control and ribosome quality-control responses to stalled or collided ribosomes [27, 37, 38]. Thus, Scp160 may act within an Asc1-associated ribosome regulatory platform that influences the fate of aggregation-prone nascent proteins such as TDP43-GFP.

The weaker phenotype of *bfr1Δ* is also informative. In our assays, deletion of *BFR1* reduced the total fraction of aggregate-containing cells but did not produce the strong shift away from severe aggregates observed in *scp160Δ* (Fig. 3). This suggests that Bfr1 contributes to the same general RNA/translation-associated pathway but is not the primary determinant of severe TDP43-GFP aggregate formation. One possibility is that Bfr1 contributes to mRNA localization or ribosome-associated RNP organization, whereas Scp160 more directly affects elongation dynamics or cotranslational handling of nascent TDP43-GFP. Thus, the distinct strengths of the two mutant phenotypes suggest related yet separable roles of Scp160 and Bfr1 in shaping the TDP43-GFP aggregation state.

Finally, although Bfr1 homologs appear largely restricted to fungi, Scp160 belongs to the conserved Vigilin family of multi-KH-domain RNA-binding proteins, whose human member is HDLBP/Vigilin [39]. Recent work has emphasized that HDLBP, like yeast Scp160, preferentially associates with ER-localized transcripts, supporting a conserved role for Vigilin-family proteins in organizing translation at the ER [40]. Thus, while the *BFR1* phenotype may reflect a fungal-specific component of this pathway, the stronger *SCP160*-dependent effect observed here raises the possibility that Vigilin-family RNA-binding proteins influence the translation-associated handling of aggregation-prone proteins more broadly. Whether HDLBP/Vigilin similarly modulates TDP-43 abundance, localization, or aggregation in mammalian cells remains an important question for future studies.

## Material and methods

### Yeast strains and culture

Wild-type *S. cerevisiae* BY4742 (MATa *his3Δ1 leu2Δ0 met15Δ0 ura3Δ0*) and the corresponding *scp160Δ* and *bfr1Δ* knockout strains were obtained from Thermo Fisher. Cultures were grown in standard rich or defined media, including YPDA (1% yeast extract, 2% peptone, 2% dextrose, supplemented with 10 mg/L adenine), YPGal (1% yeast extract, 2% peptone, 2% galactose, 1% raffinose, 10 mg/L adenine), YPRaf (1% yeast extract, 2% peptone, 2% raffinose, 10 mg/L adenine), and appropriate synthetic dropout formulations. All manipulations involving yeast cells and cultures were performed identically across samples within each experiment. Unless otherwise indicated, cultures were initiated from a single colony and grown overnight (∼16 h) at 30°C in synthetic medium lacking the appropriate amino acids, diluted into fresh medium to an OD_600_ ≈ of 0.2, and grown for an additional 3 h at 30°C prior to experimentation. Yeast transformations were done using a standard protocol [41].

### Chemicals and antibodies

Cycloheximide (CHX) was purchased from Sigma and used at concentrations of 100 μg/mL. DAPI was purchased from ThermoFisher Scientific and used at a concentration of 2.5 μg/mL. Monoclonal anti-GFP antibodies were from Santa Cruz Biotechnology (sc-9996) and used at 800 ng/mL; mouse monoclonal anti-α-tubulin antibodies (12G10) were obtained from the Developmental Studies Hybridoma Bank (DSHB; University of Iowa, Iowa City, IA, USA) and used at a 1:1000 dilution; mouse monoclonal anti-FLAG antibodies (M2) were purchased from Sigma and used at 1 μg/mL; anti-mouse IgG-HRP were from GE-Healthcare (NA931) and used at a dilution of 1:5000.

### Plasmids

TDP43-GFP expression vector is based on pRS426, and it was purchased from Addgene (#27467). Plasmids expressing Rny1-FLAG wild-type and the mutant Rny1-FLAG-W399R were described in [31].

### Cell viability and growth assays

For cell-viability assays, *WT, scp160Δ*, and *bfr1Δ* strains carrying the TDP43-GFP plasmid were grown overnight at 30°C in raffinose-containing synthetic medium lacking uracil. Cultures were diluted in the same medium to an OD_600_ ≈ of 0.2 and grown for an additional 4 h at 30°C. Cells were then divided into three subcultures, washed, adjusted to a final concentration of 2 × 10^6^ cells/ml, and six 1:5 serial dilutions were plated on glucose-containing or galactose-containing synthetic medium. Plates were incubated at 30°C for 7 days before imaging.

For growth assays, yeast cultures grown in raffinose-containing synthetic medium lacking uracil were adjusted to an OD_600_ ≈ of 0.2. Cells were pelleted, resuspended in 200 µl of raffinose-containing synthetic medium supplemented with either glucose or galactose, and inoculated into 96-well plates in triplicate. Cultures were grown for 30 h at 30°C with orbital shaking, with TDP43-GFP induction triggered during the first 4 h. OD_600_ and GFP fluorescence (ex 488 nm, em 520 nm) were measured every 30 min and automatically recorded using a BioTek Synergy HT microplate reader.

### Cell lysis and Western blot

For cell-extract preparation, cells were grown to mid-log phase at 30°C. For endpoint analyses or time-course experiments, ice-cold NaN_3_ was added to the culture to a final concentration of 100-200 mM, and 10 or 20 OD_600_ units of cells were harvested by centrifugation for 5 min at 3000 × g. Pellets were washed and resuspended in 50 or 100 µl of breaking buffer (50 mM Tris-HCl pH 7.4, 5 mM MgCl_2_) supplemented with a protease-inhibitor cocktail (Pierce). Cell suspensions were transferred to tubes containing glass beads (ø 0.25– 0.5 mm) and lysed mechanically by vortexing for a total of 10 min in 1-2 min intervals, with cooling on ice for 1 min between intervals. Cell debris was removed by centrifugation for 5 min at 1500 × g. Protein samples were denatured in 1× SDS sample buffer for 10 min at 70 °C and resolved on 12% SDS-polyacrylamide gels. Proteins were transferred to nitrocellulose membranes, detected by enhanced chemiluminescence with ECL Prime Western Blotting Detection Reagent (GE Healthcare), and imaged with an ImageQuant LAS 5000 system (Cytiva, formerly GE Healthcare).

### Sucrose-gradient centrifugation analysis

Sucrose-gradient centrifugation was carried out following the general approach described in [42]. Briefly, cultures were pre-treated with cycloheximide (100 µg/mL, 5 min), harvested by centrifugation, washed, and disrupted by glass-bead lysis. Cells were resuspended in lysis buffer (10 mM Tris-HCl, pH 7.4, 100 mM NaCl, 3 mM MgCl_2_, 100 µg/mL cycloheximide, 200 µg/mL heparin), and clarified extracts were obtained by centrifugation. Fractions corresponding to 50 A_260_ units were layered onto 11 mL 15-45% (w/v) sucrose gradients prepared in 10 mM Tris-HCl, pH 7.4, 70 mM NH_4_Cl, and 4 mM MgCl_2_. Gradients were centrifuged at 188,000 × g for 4 h 15 min at 4 °C in a Beckman SW41Ti rotor (36,000 rpm). Following centrifugation, gradients were fractionated using a Beckman recovery system coupled to a Bio-Rad EM-1 UV monitor, and A_256_ profiles were recorded digitally using Windaq software.

### Fluorescence Microscopy

Microscopic images were acquired with an Axiovert inverted microscope (Carl Zeiss Microscopy, LLC) using a 63× objective. Images were analyzed using Zen Blue software (Zeiss). For TDP43-GFP aggregate quantification, cells were imaged after 4 h of galactose induction. For each independent experiment, 100-200 cells per strain were scored manually based on GFP signal pattern as having no visible aggregates, moderate aggregates, or severe aggregates. Moderate aggregates were defined as one or more small puncta, whereas severe aggregates were defined as one large aggregate or multiple prominent aggregates within the same cell. Total aggregate-positive cells were defined as cells containing either moderate or severe aggregates.

### Statistical analyses

Statistical analyses were performed using GraphPad Prism 9. Data are shown as mean values with individual biological replicates plotted as symbols, unless otherwise indicated. For fluorescence measurements, normalized TDP43-GFP fluorescence values after 4 h of galactose induction were compared among *WT, scp160Δ*, and *bfr1Δ* strains using ordinary one-way ANOVA followed by Dunnett’s multiple-comparisons test, with *WT* used as the control group. For microscopy quantification, replicate-level percentages of total aggregate-positive cells and severe aggregate-positive cells were analyzed using ordinary one-way ANOVA followed by Tukey’s multiple-comparisons test. *P* values < 0.05 were considered statistically significant. Significance was indicated as follows: ns, not significant; \**P* < 0.05; \*\**P* < 0.01; \*\*\**P* < 0.001; \*\*\*\**P* < 0.0001.

## References

1. Soto C, and Estrada LD (2008). Protein Misfolding and Neurodegeneration. Arch Neurol. 65(2). doi: 10.1001/archneurol.2007.56.

2. Soto C (2003). Unfolding the role of protein misfolding in neurodegenerative diseases. Nat Rev Neurosci. 4(1): 49–60. doi: 10.1038/nrn1007.

3. Petroi D, Popova B, Taheri-Talesh N, Irniger S, Shahpasandzadeh H, Zweckstetter M, Outeiro TF, and Braus GH (2012). Aggregate Clearance of α-Synuclein in Saccharomyces cerevisiae Depends More on Autophagosome and Vacuole Function Than on the Proteasome. Journal of Biological Chemistry. 287(33): 27567–27579. doi: 10.1074/jbc.M112.361865.

4. Sharma N, Brandis KA, Herrera SK, Johnson BE, Vaidya T, Shrestha R, and DebBurman SK (2006). α-Synuclein Budding Yeast Model: Toxicity Enhanced by Impaired Proteasome and Oxidative Stress. JMN. 28(2): 161–178. doi: 10.1385/JMN:28:2:161.

5. Menezes R, Tenreiro S, Macedo D, Santos C, and Outeiro T (2015). From the baker to the bedside: yeast models of Parkinson’s disease. MIC. 2(8): 262–279. doi: 10.15698/mic2015.08.219.

6. Konopka CA, Locke MN, Gallagher PS, Pham N, Hart MP, Walker CJ, Gitler AD, and Gardner RG (2011). A yeast model for polyalanine-expansion aggregation and toxicity. MBoC. 22(12): 1971–1984. doi: 10.1091/mbc.e11-01-0037.

7. Hughes JN, and Thomas PQ (2013). Molecular Pathology of Polyalanine Expansion Disorders: New Perspectives from Mouse Models. In: Hatters DM, Hannan AJ, editors Tandem Repeats in Genes, Proteins, and Disease. Humana Press, Totowa, NJ; pp 135–151.

8. Johnson BS, McCaffery JM, Lindquist S, and Gitler AD (2008). A yeast TDP-43 proteinopathy model: Exploring the molecular determinants of TDP-43 aggregation and cellular toxicity. Proc Natl Acad Sci USA. 105(17): 6439–6444. doi: 10.1073/pnas.0802082105.

9. Stella R, Bertoli A, Lopreiato R, and Peggion C (2025). A Twist in Yeast: New Perspectives for Studying TDP-43 Proteinopathies in S. cerevisiae. JoF. 11(3): 188. doi: 10.3390/jof11030188.

10. Lindström M, and Liu B (2018). Yeast as a Model to Unravel Mechanisms Behind FUS Toxicity in Amyotrophic Lateral Sclerosis. Front Mol Neurosci. 11: 218. doi: 10.3389/fnmol.2018.00218.

11. Gendron TF, Josephs KA, and Petrucelli L (2010). Review: transactive response DNAbinding protein 43 (TDP-43): mechanisms of neurodegeneration. Neuropathol Appl Neurobiol. 36(2): 97–112. doi: 10.1111/j.1365-2990.2010.01060.x.

12. Ratti A, and Buratti E (2016). Physiological functions and pathobiology of TDP-43 and FUS/TLS proteins. J Neurochem. 138 Suppl 1: 95–111. doi: 10.1111/jnc.13625.

13. Guo L, and Shorter J (2017). Biology and Pathobiology of TDP-43 and Emergent Therapeutic Strategies. Cold Spring Harb Perspect Med. 7(9): a024554. doi: 10.1101/cshperspect.a024554.

14. Prasad A, Bharathi V, Sivalingam V, Girdhar A, and Patel BK (2019). Molecular Mechanisms of TDP-43 Misfolding and Pathology in Amyotrophic Lateral Sclerosis. Front Mol Neurosci. 12: 25. doi: 10.3389/fnmol.2019.00025.

15. Suk TR, and Rousseaux MWC (2020). The role of TDP-43 mislocalization in amyotrophic lateral sclerosis. Mol Neurodegeneration. 15(1): 45. doi: 10.1186/s13024-020-00397-1.

16. Johnson BS, Snead D, Lee JJ, McCaffery JM, Shorter J, and Gitler AD (2009). TDP-43 is intrinsically aggregation-prone, and amyotrophic lateral sclerosis-linked mutations accelerate aggregation and increase toxicity. J Biol Chem. 284(30): 20329–20339. doi: 10.1074/jbc.M109.010264.

17. Armakola M, Higgins MJ, Figley MD, Barmada SJ, Scarborough EA, Diaz Z, Fang X, Shorter J, Krogan NJ, Finkbeiner S, Farese RV, and Gitler AD (2012). Inhibition of RNA lariat debranching enzyme suppresses TDP-43 toxicity in ALS disease models. Nat Genet. 44(12): 1302–1309. doi: 10.1038/ng.2434.

18. Elden AC, Kim H-J, Hart MP, Chen-Plotkin AS, Johnson BS, Fang X, Armakola M, Geser F, Greene R, Lu MM, Padmanabhan A, Clay-Falcone D, McCluskey L, Elman L, Juhr D, Gruber PJ, Rüb U, Auburger G, Trojanowski JQ, Lee VM-Y, Van Deerlin VM, Bonini NM, and Gitler AD (2010). Ataxin-2 intermediate-length polyglutamine expansions are associated with increased risk for ALS. Nature. 466(7310): 1069–1075. doi: 10.1038/nature09320.

19. Ju S, Tardiff DF, Han H, Divya K, Zhong Q, Maquat LE, Bosco DA, Hayward LJ, Brown RH, Lindquist S, Ringe D, and Petsko GA (2011). A Yeast Model of FUS/TLS-Dependent Cytotoxicity. PLoS Biol. 9(4): e1001052. doi: 10.1371/journal.pbio.1001052.

20. Sun Z, Diaz Z, Fang X, Hart MP, Chesi A, Shorter J, and Gitler AD (2011). Molecular Determinants and Genetic Modifiers of Aggregation and Toxicity for the ALS Disease Protein FUS/TLS. PLoS Biol. 9(4): e1000614. doi: 10.1371/journal.pbio.1000614.

21. Frey S, Pool M, and Seedorf M (2001). Scp160p, an RNA-binding, polysome-associated protein, localizes to the endoplasmic reticulum of Saccharomyces cerevisiae in a microtubule-dependent manner. J Biol Chem. 276(19): 15905–15912. doi: 10.1074/jbc.M009430200.

22. Hirschmann WD, Westendorf H, Mayer A, Cannarozzi G, Cramer P, and Jansen R-P (2014). Scp160p is required for translational efficiency of codon-optimized mRNAs in yeast. Nucleic Acids Research. 42(6): 4043–4055. doi: 10.1093/nar/gkt1392.

23. Hogan DJ, Riordan DP, Gerber AP, Herschlag D, and Brown PO (2008). Diverse RNA-binding proteins interact with functionally related sets of RNAs, suggesting an extensive regulatory system. PLoS Biol. 6(10): e255. doi: 10.1371/journal.pbio.0060255.

24. Lang BD (2001). The brefeldin A resistance protein Bfr1p is a component of polyribosome-associated mRNP complexes in yeast. Nucleic Acids Research. 29(12): 2567–2574. doi: 10.1093/nar/29.12.2567.

25. Castells-Ballester J, Rinis N, Kotan I, Gal L, Bausewein D, Kats I, Zatorska E, Kramer G, Bukau B, Schuldiner M, and Strahl S (2019). Translational Regulation of Pmt1 and Pmt2 by Bfr1 Affects Unfolded Protein O-Mannosylation. Int J Mol Sci. 20(24): 6220. doi: 10.3390/ijms20246220.

26. Manchalu S, Mittal N, Spang A, and Jansen R-P (2019). Local translation of yeast ERG4 mRNA at the endoplasmic reticulum requires the brefeldin A resistance protein Bfr1. RNA. 25(12): 1661–1672. doi: 10.1261/rna.072017.119.

27. Baum S, Bittins M, Frey S, and Seedorf M (2004). Asc1p, a WD40-domain containing adaptor protein, is required for the interaction of the RNA-binding protein Scp160p with polysomes. Biochemical Journal. 380(3): 823–830. doi: 10.1042/bj20031962.

28. Sezen B, Seedorf M, and Schiebel E (2009). The SESA network links duplication of the yeast centrosome with the protein translation machinery. Genes Dev. 23(13): 1559–1570. doi: 10.1101/gad.524209.

29. Weidner J, Wang C, Prescianotto-Baschong C, Estrada AF, and Spang A (2014). The polysome-associated proteins Scp160 and Bfr1 prevent P body formation under normal growth conditions. Journal of Cell Science. jcs.142083. doi: 10.1242/jcs.142083.

30. Lapointe CP, Wilinski D, Saunders HAJ, and Wickens M (2015). Protein-RNA networks revealed through covalent RNA marks. Nat Methods. 12(12): 1163–1170. doi: 10.1038/nmeth.3651.

31. Shcherbik N (2013). Golgi-mediated glycosylation determines residency of the T2 RNase Rny1p in Saccharomyces cerevisiae. Traffic. 14(12): 1209–1227. doi: 10.1111/tra.12122.

32. Bharathi V, Bajpai A, Parappuram IT, and Patel BK (2022). Elevated constitutive expression of Hsp40 chaperone Sis1 reduces TDP-43 aggregation-induced oxidative stress in Ire1 pathway dependent-manner in yeast TDP-43 proteinopathy model of amyotrophic lateral sclerosis. Biochemical and Biophysical Research Communications. 595: 28–34. doi: 10.1016/j.bbrc.2022.01.073.

33. Stagoj MN, Comino A, and Komel R (2005). Fluorescence based assay of GAL system in yeast Saccharomyces cerevisiae. FEMS Microbiol Lett. 244(1): 105–110. doi: 10.1016/j.femsle.2005.01.041.

34. Collart MA, and Weiss B (2020). Ribosome pausing, a dangerous necessity for co-translational events. Nucleic Acids Research. 48(3): 1043–1055. doi: 10.1093/nar/gkz763.

35. Cheng MHK, Hoffmann PC, Franz-Wachtel M, Sparn C, Seng C, Maček B, and Jansen R-P (2018). The RNA-Binding Protein Scp160p Facilitates Aggregation of Many Endogenous Q/N-Rich Proteins. Cell Rep. 24(1): 20–26. doi: 10.1016/j.celrep.2018.06.015.

36. Wen J-H, He X-H, Feng Z-S, Li D-Y, Tang J-X, and Liu H-F (2023). Cellular Protein Aggregates: Formation, Biological Effects, and Ways of Elimination. Int J Mol Sci. 24(10): 8593. doi: 10.3390/ijms24108593.

37. Ikeuchi K, Tesina P, Matsuo Y, Sugiyama T, Cheng J, Saeki Y, Tanaka K, Becker T, Beckmann R, and Inada T (2019). Collided ribosomes form a unique structural interface to induce Hel2-driven quality control pathways. EMBO J. 38(5): e100276. doi: 10.15252/embj.2018100276.

38. Kuroha K, Zinoviev A, Hellen CUT, and Pestova TV (2018). Release of Ubiquitinated and Non-ubiquitinated Nascent Chains from Stalled Mammalian Ribosomal Complexes by ANKZF1 and Ptrh1. Mol Cell. 72(2): 286-302.e8. doi: 10.1016/j.molcel.2018.08.022.

39. Feicht J, and Jansen R-P (2024). The high-density lipoprotein binding protein HDLBP is an unusual RNA-binding protein with multiple roles in cancer and disease. RNA Biology. 21(1): 312–321. doi: 10.1080/15476286.2024.2313881.

40. Zinnall U, Milek M, Minia I, Vieira-Vieira CH, Müller S, Mastrobuoni G, Hazapis O-G, Del Giudice S, Schwefel D, Bley N, Voigt F, Chao JA, Kempa S, Hüttelmaier S, Selbach M, and Landthaler M (2022). HDLBP binds ER-targeted mRNAs by multivalent interactions to promote protein synthesis of transmembrane and secreted proteins. Nat Commun. 13(1): 2727. doi: 10.1038/s41467-022-30322-7.

41. Gietz RD, and Schiestl RH (2007). Large-scale high-efficiency yeast transformation using the LiAc/SS carrier DNA/PEG method. Nat Protoc. 2(1): 38–41. doi: 10.1038/nprot.2007.15.

42. Shcherbik N, Chernova TA, Chernoff YO, and Pestov DG (2016). Distinct types of translation termination generate substrates for ribosome-associated quality control. Nucleic Acids Res. 44(14): 6840–6852. doi: 10.1093/nar/gkw566.

